# Overlapping *oriC* and centromere-like functions in secondary genome replicons determine their maintenance independent of chromosome I in *Deinococcus radiodurans*

**DOI:** 10.1101/2020.03.28.013953

**Authors:** Ganesh K Maurya, Hari S. Misra

**Affiliations:** Molecular Biology Division, Bhabha Atomic Research Centre, Mumbai, India – 400085; Homi Bhabha National Institute, Mumbai, India – 400094

## Abstract

The *Deinococcus radiodurans* multipartite genome system (MGS) consists of chromosome I (ChrI) and secondary genome elements; Chr II and megaplasmid (MP). The sequences upstream to *parAB* operons in Chr II (*cisII*) and MP *(cisMP*) helped an *E. coli* plasmid maintenance in *D. radiodurans* and showed sequence specific interactions with DnaA and ParBs. The cells devoid of *cisII* (Δ*cisII*) or *cisMP* (Δ*cisMP*) showed reduced γ radiation resistance and copy number of Chr II and MP. <u>F</u>luorescent <u>R</u>eporter-<u>O</u>perator System (FROS) developed for ChrI, ChrII and MP in Δ*cisII* or Δ*cisMP* mutants showed no change in wild type pattern of Chr I localization. However, the relative copy numbers of Chr II and MP had reduced while anucleate cells had increased in mutants. These results suggested that *cisII* and *cisMP* elements contain both *ori* and centromere-like functions, and like other MGS bacteria, the Chr I and secondary genome are maintained independently in *D. radiodurans*.

## 1. Introduction

Genome duplication and segregation are highly coordinated macromolecular events critical for growth of any organism^1^. The chromosomal replication starts at *oriC* a well conserved region comprised of the non-palindromic repeats of 9 mer as DnaA boxes and 13 mer AT rich repeats^2–5^. In the best studied, single circular chromosome harbouring, *E. coli* model, the DnaA boxes are bound by ATP-complexed DnaA protein following the recruitment of replication initiation complex comprised of DnaB helicase with the help of DnaC, primase and DNA polymerase III ^6,7^. Segregation of duplicated replicons is also linked with the number of copies per cell. Various models of active genome segregation by tripartite genome segregation (TGS) system have been suggested for single circular chromosome and limited copy large plasmids^8^. In TGS, the nucleation of adapter protein like ParB on centromeric sequences is a limiting step for faithful segregation of duplicated chromosome^9^. The distribution of high copy number plasmids in dividing population is believed to be a random passive process ^10^. It has been observed that the initiation of DNA replication and segregation of duplicated genome occurs concurrently and therefore believed to be interdependently regulated in bacteria ^11^. In majority of the cases where DNA replication and segregation have been studied, the centromeric sequences are found close to *ori*. For instance, *B. subtilis* and *crescentus*, the primary *parS* sites are located with 8-10 kb from *oriC* ^12^. Likewise, the segregation proteins interact directly with replication initiation protein DnaA in many bacteria ^11,13^. The organisation and dynamics of nucleoid during growth have been studied using genetic, microscopy and chromosome conformation capture techniques in rod- or crescent-shaped bacteria like *E. coli*, *B. subtilis* and *C. crescentus* ^1, 14–17^ and in cocci like *Streptococcus pneumonia* ^18^ and *Staphylococcus aureus* ^19,20^.

Recently, many bacteria have been discovered with multiple copies of multipartite genome system comprising of more than one chromosome and megaplasmids ^21^. Studies on genome maintenance in bacteria harbouring multipartite genome (MGH) are limited to a few examples like *V. cholerae*, *Burkholderia cenocepacia* and *Rhodobacter sphearoides* ^22–24^. In these bacteria, the partitioning components of primary chromosome are phylogenetically similar with the chromosome of single circular chromosome harbouring bacteria but are different from the elements encoded on secondary genome elements. Since, the sizes of the secondary genome elements vary, the synchronization in their duplication and segregation with cytokinetic events of cell division is quite intriguing. However, the real time monitoring of DNA replication in *V. cholerae* harbouring multiple size genome elements have revealed staggered replication initiation for different elements to allow the replication termination roughly around the same time ^25,26^. The nucleoids dynamics during genome segregation have also been studied in some MGH bacteria like *V. cholerae, R. sphaeroides, Shinorhizobium meliloti* and *D. radiodurans* ^22, 24, 27,28^.

*D. radiodurans*, a Gram-positive non-pathogenic bacterium characterized for extraordinary radioresistance, also harbours multipartite genome system comprised of chromosome I (2,648,638bp), chromosome II (412,348bp) and megaplasmid (177,466bp) ^29^. Earlier, Minsky and colleagues demonstrated that the polyploid multipartite genome of *D. radiodurans* is packaged in form of a doughnut shape toroidal structure, which was implicated to its extreme resistance for DNA double strand breaks (DSBs) and efficient DSB repair. The toroidal packaging of polyploid multipartite genome raises the mechanistic questions on the “*oriC*” regulation of chromosomal replication, segregation of genome elements and DSB repair and recovery of individual genome elements identity post DNA breaks. Recently, the nucleoid dynamics in *D. radiodurans* has been studied using high resolution imaging, which reports that deinococcal nucleoid is highly condensed but remain dynamic to adapt multiple distinct configurations during cell cycle progression ^28^. They studied chromosome I (Chr I) dynamics during different stages of cell cycle and showed that the *oriC* in Chr I remains radially distributed around the centrally positioned *ter* (terminus) sites. The characterization of *oriC* and centromere-like sequences in chromosome II (Chr II) and megaplasmid (MP), and their localisation are untraced in the nucleoid of *D. radiodurans.* Here, we report that the direct repeats located upstream of *parAB* operons in Chr II (*cisII*) and MP (*cisM*P) contain both origin of replication (hereafter referred ‘*ori*’) centromere-like functions. We demonstrated that both DnaA interacts with both these *cis* elements while ParBs interact with cognate *cis* elements *in vitro,* and deletion of these *cis* elements leads to drastic reduction in copy numbers of respective replicons and partial loss of radioresistance of this bacterium. These deletion mutants also showed the increased frequency of cells devoid of cognate replicons. Thus, we identified ‘*ori’* and centromere-like functions in these *cis* elements encoded on ChrII and MP in *D. radiodurans.* Using a *tetO*-TetR-GFP based <u>F</u>luorescent <u>R</u>eporter <u>O</u>perator System (FROS) approach, the ‘*ori’* in Chr I, Chr II and MP were marked, and showed that Chr I localization and dynamics is independent of secondary genome replicons (Chr II and MP) in this multipartite genome harboring bacterium.

## 2. Material and Methods

### 2.1 Bacterial strains, plasmids and materials

All the oligonucleotides used in this study are listed in Table S1 and all bacterial strains and plasmids are listed in Table S2. *D. radiodurans* R1 (ATCC13939) was grown in TGY (1% Tryptone, 0.1% Glucose and 0.5% Yeast extract) medium at 32°C. *E. coli* strain NovaBlue was used for cloning and maintenance of all the plasmids while *E. coli* strain BL21 (DE3) pLysS was used for the expression of recombinant proteins. The pNOKOUT plasmid (a suicide plasmid in *D. radiodurans*) was used for generating gene knockouts and for bioassay of *cis* elements as origin of replication in *D. radiodurans*. Standard protocols for all recombinant techniques were used as described earlier ^30^. Molecular biology grade chemicals and enzymes were procured from Merck Inc. and New England Biolabs, USA. Radiolabeled nucleotides were purchased from Board of Radiation and Isotope Technology, Department of Atomic Energy, India (BRIT, India).

### 2.2 Bioinformatic analysis

The nucleotide sequences close to the origin region in the circular map of Chr II and MP were analysed for potential repeats and consensus sequence using Mellina II web tool ^31^ and WebLogo online tool ^32^ and tentatively named as *cisII and cisMP,* respectively. The putative DnaA boxes in *cisII* were predicted using DOriC database (DoriC accession number – ORI10030006) ^33^. The DnaA boxes in *cisMP* were marked based on sequence homology with known DnaA boxes from other bacteria.

### 2.3 Cloning of *cis* elements and their maintenance

The *cisII* and *cisMP* were PCR amplified from genomic DNA of *D. radiodurans* using cisIIFw & cisIIRw and cisMPFw & cisMPRw primers, respectively (Table S1). The *cisII* was cloned at *Xba*I while *cisMP* at *Apa*I and *Eco*RI sites in pNOKOUT ^34^ to yield pNOKcisII and pNOKcisMP plasmids, respectively (Table S2). These plasmids were transformed in both wild type and *recA* mutant of *D. radiodurans* and transformants were scored at kanamycin (6μg/ml) containing TYG agar plates. The bacterial growth was monitored at O.D. 600 nm, in the presence of required selection pressure at 32°C for 24h using Synergy H1 Hybrid Multi-mode microplate reader, Bio-Tek. The maintenance of pNOKcisII and pNOKcisMP plasmids in *D.* radiodurans was confirmed with antibiotic resistance as well as by isolating these plasmids from transformant as described earlier ^35^. The plasmid DNA was digested with *Xba*I in case of pNOKcisII and *Apa*I and *Eco*RI in case of pNOKcisMP and release of desired inserts was verified on agarose gel.

### 2.4 Protein purification

The recombinant DnaA of *D. radiodurans* (hereafter referred as DnaA unless other mentioned) expressing on pETDnaA in E. coli BL21 (DE3) pLysS was purified by Ni-NTA affinity chromatography, as described earlier ^36^. In brief, E. coli cells were treated with 0.5 mM isopropyl-β-D-thio-galacto-pyranoside (IPTG) and cell pellet was suspended in buffer A (20 mM Tris–HCl, pH 7.6, 300 mM NaCl and 10% glycerol), 0.5 mg/ml lysozyme, 1 mM PMSF, 1mM MgCl_2_, 0.05 % NP-40, 0.05 % TritonX-100, protease inhibitor cocktail), and incubated on ice for 1h. The cell lysate made by sonication was centrifuged at 11,000 rpm for 30 min at 4^◦^C and the clear supernatant was purified through Ni-NTA column chromatography (GE Healthcare) following kit protocols. The column was thoroughly washed with buffer A containing 50 mM imidazole. Recombinant protein was eluted in steps using 100 mM, 200 mM, 250 mM and 300 mM imidazole in buffer A and analyzed on 10% SDS-PAGE. Fractions with >95% pure protein were pooled and re-purified from HiTrap Heparin HP affinity columns (GE Healthcare Life sciences) using a linear gradient of 0 – 300 mM NaCl, followed by precipitation with 30% w/v ammonium sulphate at 8°C and molecular exclusion chromatography using standard protocol. The recombinant ParBs encoded on chromosome II (ParB2) and megaplasmid (ParB3) in *D. radiodurans* were also purified from recombinant *E. coli* using Ni-NTA affinity chromatography followed by molecular exclusion chromatography as described earlier ^37^. Protein concentration was determined by taking OD at 280 nm in Nano drop (Synergy H1 Hybrid Multi-Mode Reader Biotek) and mass extinction co-efficient of the proteins.

### 2.5 DNA-protein interaction studies

DNA binding activity of DnaA, ParB2 and ParB3 was monitored using electrophoretic mobility shift assay (EMSA) with different version of *cis* elements as DNA substrates ^36^. In brief, full length *cisII* and *cisMP* and their repeat derivatives (Fig. S2) were PCR amplified using sequence-specific primers (Table S1). The PCR products were gel purified and labeled with [γ-^32^P] ATP using T4 polynucleotide kinase. Two repeats and one repeat of both the *cis* elements were chemically synthesised, radiolabelled with [γ-^32^P] ATP using T4 polynucleotide kinase, and annealed with unlabelled complimentary strands (Table S1). These substrates were incubated with different concentrations of purified recombinant DnaA or ParB proteins in a reaction buffer containing 50 mM Tris-HCl (pH 8.0), 75mM KCl, 5mM MgSO_4_ and 0.1mM DTT at 37°C for 15 min. For the competition assay, saturating ratios of DnaA or ParBs to DNA substrates were chased with different concentration of non-specific DNA (NS-DNA) of similar length (Table S1) and a 10-fold higher concentration of same *cis* sequence. Mixtures were separated on 6-10 % native PAGE gel as required, dried, and autoradiogram was developed on X-ray films. The band intensity of bound and unbound fraction was determined by using Image J 2.0 software. The fraction of DNA probe bound to the protein was plotted as a function of the protein concentration by using GraphPad Prism 5. The Kd for the curve fitting of individual plots was determined as described before ^36,38^.

### 2.6 Construction of deletion mutants and cell survival studies

The constructs for generating knockout of both the *cis* elements were made in pNOKOUT plasmid as described earlier ^34^. In brief, ~500 bps upstream and ~500 bps downstream fragments to the *cisII* and *cisMP* were PCR amplified with the primers (Table S1) and cloned at *Kpn*I & *Eco*RI, and *Bam*HI & *Xba*I sites in pNOKOUT plasmid to yielded pNOKCII and pNOKCM plasmids, respectively (Table S2). These plasmids were linearized with *Xmn*I and transformed into *D. radiodurans* cells. The homozygous replacement of *cisII* and *cisMP* by *nptII* was achieved through several rounds of sub-culturing and ascertained by diagnostic PCR using flanking primers (Table S1) of the target as described earlier (Maurya et al., 2018). The confirmed mutant cells were maintained in the absence of selection pressure but ensured for antibiotic resistance before the experiments.

The survival of Δ*cisII* and Δ*cisMP* deletion mutants was monitored under normal conditions and in response to different doses of gamma radiation as described earlier ^39^. In brief, the cells were grown in the absence and presence of antibiotics and exposed to different doses of γ-radiation (2, 4, 6 and 9 kGy) at dose rate of 1.81 kGy/h using Gamma Cell 5000 irradiator (^60^Co; Board of Radiation and Isotopes Technology, DAE, India). The equal number of irradiated cells and their SHAM controls were washed in PBS and the different dilutions were plated on TYG agar plates with antibiotics as required, in replicates. These plates were incubated at 32°C for 45-48 h and colony forming units (CFU) were counted.

### 2.7 Estimation of genome copy number

The genome copy number of each replicon in *D. radiodurans* was determined in exponentially growing cells. The density and cell numbers of independent sample were normalized for a fix OD 600 nm and by cfu. These cells were harvested, washed with 70 % ethanol solution and re-suspended in a lysis solution containing 10mM Tris pH 7.6, 1 mM EDTA and 4 mg/ml lysozyme. The mixtures were incubated at 37°C for 1 h, lysed by repeated cycles of freezing and thawing and centrifuged at 10000 rpm for 5 min. An aliquot of clear supernatant was serially diluted, and 0.1 ml was used for genome copy number determination using quantitative Real-Time PCR as described in ^36,40^. In brief, a fragment of about 300 bps was PCR amplified and purified using Gel Extraction kit. The amount of DNA was quantified by nanodrop and the DNA concentrations were calculated using the molecular mass computed with ‘oligo calc’ (www.basic.northwestern.edu/biotools). A serial dilution was made for each standard fragment and qPCR was carried out by following the MIQE guidelines using Roche Light cycler ^41^ and the cycle threshold (Ct) values were determined. The qPCR analysis involves three different genes from each genome replicon of *D. radiodurans* with similar PCR efficiency. For instance, *ftsE, ftsK* and *ftsZ* for chromosome I, DR_A0155, DR_A0182 and *pprA* for chromosome II, DR_B0003, DR_B0030 and DR_B0104 for megaplasmid and DR_C0001 and DR_C0018 and DR_C0034 for small plasmid (Table S1). The experiments were carried out with three independent biologic replicates for each sample. The replicon copy number is quantified by comparing the results with a standard plot. The copy number of each replicon as determined by considering the amplification of all genes was calculated using the cell number present at the time of cell lysis. Average of copy number reflected from all three genes per replicon was analysed using statistical analysis. The copy numbers of pNOKcisII and pNOKcisMP in *D. radiodurans* were determined using similar approach as described above except that the genes used were *nptII* (kanamycin resistance gene) and *bla* (ampicillin resistance gene) from plasmid backbone (Table S1).

### 2.8 Construction of *tetO* insertion derivatives of genome elements in *D. radiodurans*

We used *tetO* operator cassette (an array of 240 repeats) from pLAU44 ^42^. The whole plasmid containing operator sequences was integrated near the origin of Chr I, Chr II and MP in *D. radiodurans.* In brief, we PCR amplified the region (10713-11715) corresponding to 1.5° of Chr I using ChrI(1.5°)Fw and ChrI(1.5°)Rw primers, region (4659-c5691) corresponding to 4° of Chr II using ChrII(4°)Fw and ChrII(4°)Rw primers, and region (2203-3000) that corresponds to 4.5° of MP using Mp(4.5°)Fw and Mp(4.5°)Rw. These PCR fragments were cloned in pLAU44 at *Xba*I-*Sca*I sites to yield p44Ch1, p44Ch2 and p44MP plasmids having homologous sequences of Chr I, Chr II and MP, respectively. Further, an expressing cassette of spectinomycin resistance gene was amplified from pVHS559 ^38^ using SpecFw and SpecRw primers and cloned at *Nhe*I-*Xho*I sites in p44Ch1, p44Ch2 and p44MP to give p44SCh1, p44SCh2 and p44SMP, respectively. These constructs were confirmed by restriction digestion, PCR and DNA sequencing.

For integration of *tetO* in genome replicons, the wild type *D. radiodurans* and its Δ*cisII* and Δ*cisMP* mutant were separately transformed with p44SCh1, p44SCh2, p44SMP plasmids and transformants were scored under antibiotic selection. The homogenous insertion of *tetO* repeats in genome was achieved through several rounds of sub-culturing and ascertained by diagnostic PCR using different sets of primers (Table S1). The resultant strains were named with suffix as R1::ChrI-*tetO,* R1::ChrII-*tetO*, R1::MP-*tetO,* Δ*cisII*::ChrI-*tetO,* Δ*cisII*::ChrII-*tetO*, Δ*cisII*::MP-*tetO,* Δ*cisMP*::ChrI-*tetO,* Δ*cisMP*::ChrII-*tetO* and Δ*cisMP*::MP-*tetO* in *D. radiodurans* and listed in Table S2.

### 2.9 Construction of TetR-GFP expression plasmid in *D. radiodurans*

The reporter gene *tetR* was PCR amplified from pLAU53 ^42^ using gene specific primers (Table S1) and cloned at *Sac*I and *Sal*I sites in pDSW208 ^43^ to give pDTRGFP plasmid. Further, we PCR amplified *tetR-gfp* fusion fragment from pDTRGFP plasmid using TetRscIApIFw and GFPXbaIRw primers and cloned in pRADgro ^39^ at *Apa*I-*Xba*I sites to give pRADTRGFP plasmid (Table S2). The constitutive expression of TetR-GFP from their respective plasmids in *D. radiodurans* was confirmed by western blotting and fluorescence microscopy.

We transformed pRADTRGFP plasmid into *D. radiodurans* R1 and its Δ*cisII* and Δ*cisMP* mutants (Table S2) carrying *tetO* operator sequences and maintained in the presence of required antibiotics. The fluorescence microscopic studies for localization of different replicons tagged close to origin position in wild type and *cis* mutants of *D. radiodurans* were performed as described in ^36^. In brief, overnight grown cultures were harvested, washed with PBS, stained with DAPI (4’,6-diamidino-2-phenylindole, Dihydrochloride; 0.2 mg/ml) for nucleoid and Nile red (1 mg/ml) for plasma membrane. These cells were mounted on glass slides bed with 0.8% agarose and imaged under inverted fluorescence microscope (Olympus IX83) equipped with an Olympus DP80 CCD monochrome camera under DIC (Differential Interference Contrast), DAPI, FITC (Fluorescein isothiocyanate) and TRITC (Tetramethylrhodamine isothiocyanate) channels. Images were aligned and processed using in-built software ‘CellSens’. A population of 200 cells were analysed for results and data were presented as scattered plot. One representative image belonging to more than 90 % population is presented separately for DIC, DAPI, Nile Red and GFP as well as merged. Experiments were repeated independently to ensure the reproducibility and significance of these data.

## 3. Results

### 3.1 Sequences in Chr II and MP predicted as putative *‘oriC’* and centromere-like sequences

The nucleotide sequences of Chr II and MP were subjected to *in silico* search of repetitive sequences similar to known ‘*ori*’ and centromere-like sequences. Results showed 11 iterons of 18 bp with a consensus sequence T(G)CCACAAAGTGCCA(G)CAGG and GC content of 51.6%, spanning 99-544 bp region in Chr II which was named as *cisII*. Similarly, 8 iterons of 18 bp and 3 iterons of 14bp were mapped with the consensus sequence C(A)CCGCAAAGGTG(A)TCGCTA and GC content of 47%, spanning 177446-417 bp region in MP and named as *cisMP* (Fig. 1). In both *cisII* and *cisMP* elements, the repeats were mapped near their origins in circular map, and upstream to *parAB* operon. In addition, both *cisII* and *cisMP* elements also contain non-canonical putative DnaA boxes which were distinct from the 13 mer DnaA boxes (T(A/G)TA(T)TCCACA) present in the origin of replication of Chr I ^44^. Though, the DnaA boxes started just upstream of the 18 bp iterons, they extended well into the iterons resulting in a considerable overlap between both the elements, which could explain the temporal regulation of these macromolecular events by differential interaction with respective *trans* factors. In addition, both the *cis* elements contain GATC and AT rich sequences *albeit* different in numbers, which are the known signatures of “*ori*” like sequences and needed for replication initiation in bacteria (Fig 1). In a previous study, 3 centromeric-like regions with consensus of GTTTC(A)G(A)C(A)GT (C)GG(T)A(G)AAC were mapped at 3.5° (*segS*1), 72.9° (*segS*2) and 231.8° (*segS*3) in Chr I ^38^. The predicted centromere-like sequences in Chr II and MP are found to be distinct from the conserved *segS* sequence of Chr I. Thus, we found that *cisII* and *cisMP* elements have DnaA boxes as ‘*oriC’* feature and the iterons as centromere-like sequences that may contribute in genome segregation. The presence of distinct ‘*ori*’ and centromeric signatures in the Chr I and secondary genome elements clearly indicated a strong possibility of their independent maintenance in this bacterium as reported in other MGH bacteria like *B. cenocepacia* and *V. cholerae* ^12, 23, 45^.

**Figure 1:**
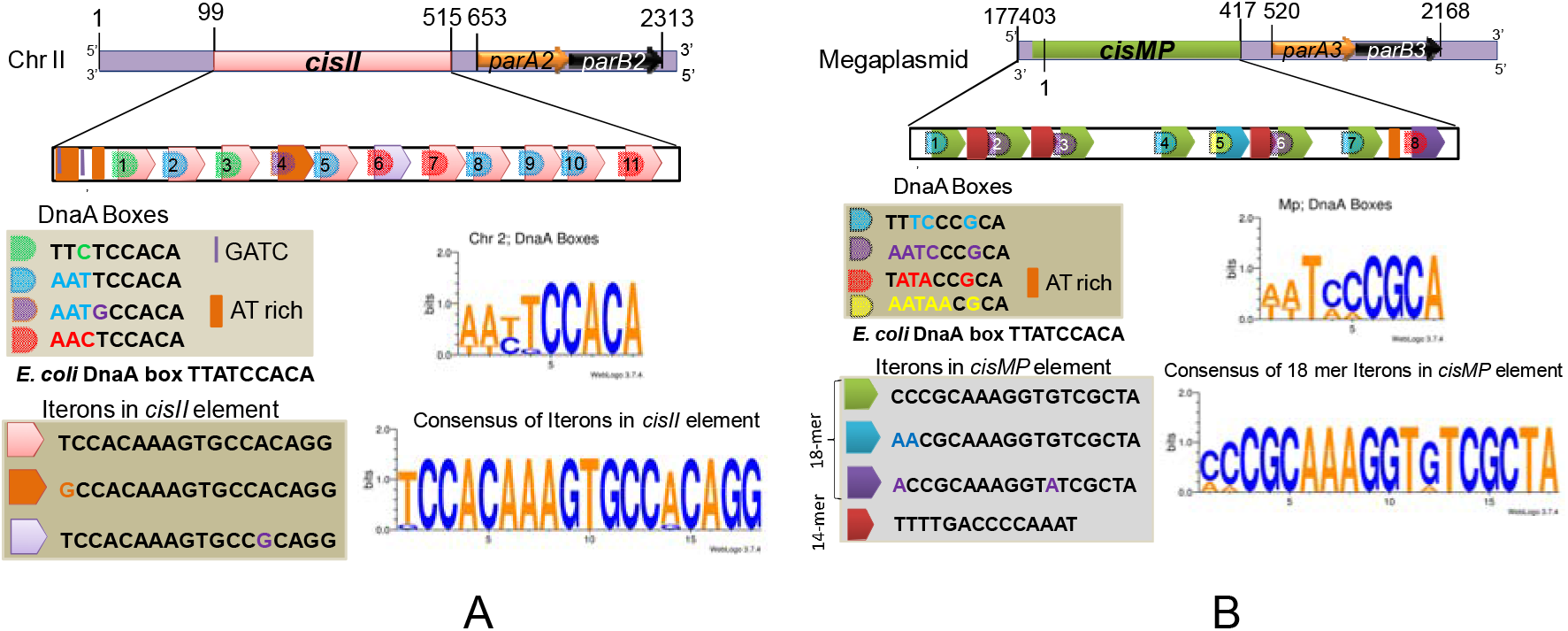
Schematic presentation of direct repeats located upstream to *parAB* operon in chromosome II (*cisII*) and megaplasmid (*cisMP*). The 99-554 bp region of chromosome II (A) and 177446 to 417bp region of megaplasmid (B) were analysed *in silico* for the structure of direct repeats and their sequence compositions of DnaA boxes and iterons.

### 3.2 *cisII* and *cisMP* elements confer *ori* function in *D. radiodurans*

Since, *cisII* and *cisMP* sequences contain typical DnaA boxes and are located upstream to *parAB* operons, a possibility of these elements acting as origin of replication and centromere-like elements in *D. radiodurans* was investigated. We used pNOKOUT a *colE1* based *E. coli* plasmid for stability assay in *D. radiodurans*. For that, these elements were separately cloned in pNOKOUT and the resultant plasmids pNOKCisII and pNOKCisMP were transformed into *D. radiodurans*. Unlike pNOKOUT vector, the transformants harbouring pNOKCisII or pNOKCisMP were able to grow under kanamycin (6 μg/ml) selection both on TGY plates and in broth, which was found to be nearly similar to wild type grown without antibiotics (Fig. 2A, C). In order to rule out that the expression of plasmid born antibiotic resistance was not due to integration of these plasmids into chromosome, the maintenance of these plasmids was checked in recombinant negative Δ*recA* mutant of *D. radiodurans*. The results were same as wild type and the Δ*recA* transformants of pNOKCisII or pNOKCisMP plasmids could grow in the presence of required dose of antibiotic (Fig. 2B). Further, the independent multiplication of these plasmids was confirmed independently by isolating them from kanamycin resistant recombinant *D. radiodurans* and restriction analysis. Results showed the release of expected size insert from recombinant derivatives of pNOKOUT plasmid (Fig. 2D). When we measured the copy number by qPCR, it was 6.86 ± 0.71 for pNOKCisII and 12.15 ± 0.92 for pNOKCisMP, which was found to be very close to *in vivo* copy number of Chr II and MP, respectively in *D. radiodurans*. These results together provided the clear evidence that pNOKCisII and pNOKCisMP plasmids were independently maintained in *D. radiodurans* because of the presence of *cisII* and *cisMP* elements, as the vector without them could not survive, suggesting the ‘*ori*’ nature of *cisII* and *cisMP* elements for cognate replicons in this bacterium.

**Figure 2.**
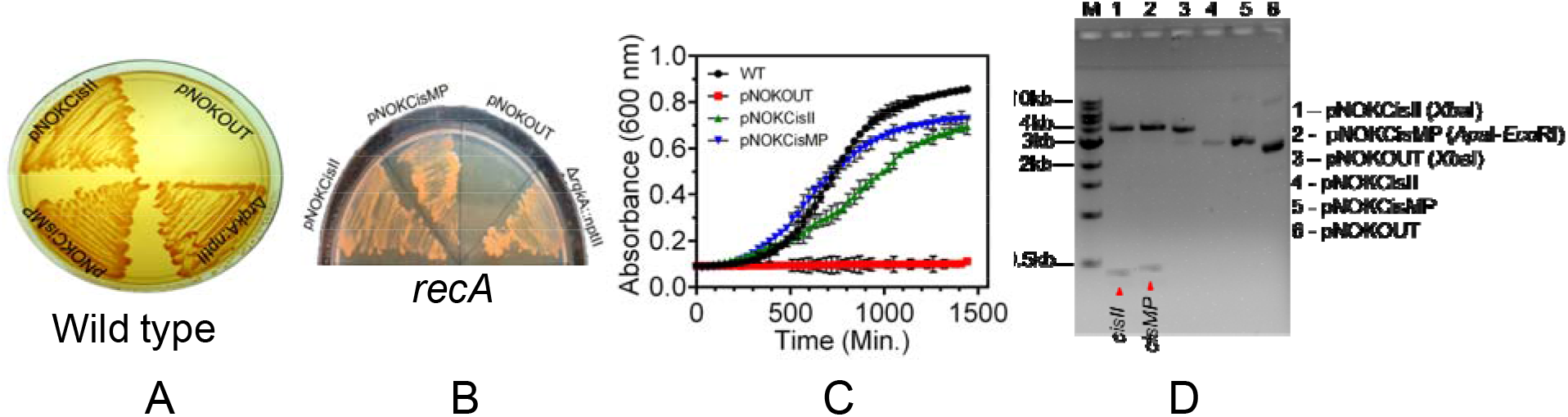
Role of *cisII* and *cisMP* elements in the replication of *Escherichia coli* plasmid into *Deinococcus radiodurans*. The *cisII* and *cisMP* elements were cloned in *E. coli* cloning vector pNOKOUT, and the resultant plasmids pNOKcisII and pNOKcisMP were transformed into wild type (A) and *recA* mutant (B). These transformants were grown in the presence of kanamycin (6 μg / ml) and growth was monitored on TGY agar plate with *rqkA* deletion mutant as positive control and vector as negative control. The growth characteristic of these transformants grown in TGY broth supplemented with antibiotic was compared with wild type cells grown in TGY broth without antibiotics (C). The plasmids isolated from recombinant *D. radiodurans* cells growing in the presence of antibiotic were digested with restriction enzymes and the release of *cisII* and *cisMP* fragments was analysed on agarose gel and compared with empty vector (D).

### 3.3 *cisII* and *cisMP* elements provide sequence specific interaction to DnaA

To understand the molecular mechanisms of *cisII* and *cisMP* roles as *ori*, the recombinant deinococcal replication initiator protein DnaA was purified (Fig. S1), and its DNA binding activity was checked with radiolabelled *cisII* and *cisMP* elements and their repeat derivatives (Fig. S2). Results showed that DnaA establish the sequence specific interaction with both *cisII* and *cisMP* albeit with different affinity (Fig. 3) and its affinity varies with the number of these repeats. For instance, DnaA affinity to *cisII* was higher than *cisMP*. The affinity of DnaA to both the *cis* elements had reduced with the reduction of repeats’ number (Fig. 3, Table 1). As expected, the DnaA bound to either of *cis* elements remained unaffected even in the presence of a 100-fold higher molar concentration of non-specific DNA, which could be titrated out completely with 10-fold molar excess of specific DNA (Fig. 3). The formation of DnaA complex with *cisII* and *cisMP* containing several directs repeats clearly suggests the ‘*ori*’ nature of both the *cis* elements in *D. radiodurans*.

**Figure 3.**
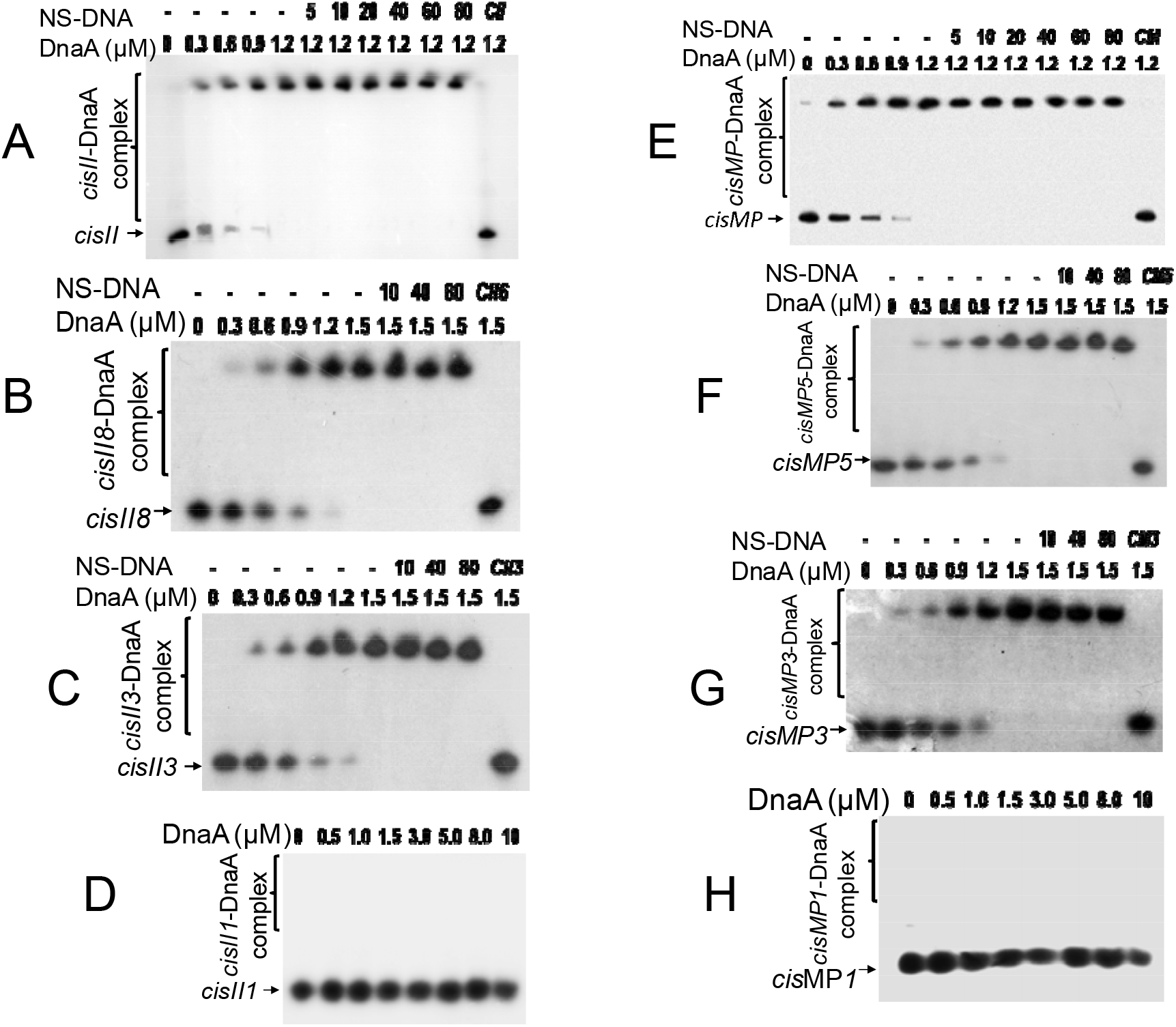
DnaA interaction with *cisII* and *cisMP* and their repeats variants. The radiolabelled full length *cisII* containing 11 repeats (A), 8 repeats (B), 3 repeats (C) and 1 repeat (D) (Fig S2A) were incubated with the increasing concentration of DnaA. Similarly, the radiolabelled full length *cisMP* containing 8 repeats (E), 5 repeats (F), 3 repeats (G) and single repeat (H) were incubated with the increasing concentration of DnaA. A saturating concentration of DnaA-DNA ratio was chased with increasing molar ratio of non-specific DNA (NS-DNA) and products were analysed on native PAGE. The fraction of DNA bound with proteins was quantified densitometrically and plotted as a function of protein concentration using Graphpad PRISM. The Kd for the curve fitting of individual plots was determined and given in Table 1.

**Table 1.** The dissociation constant (Kd) value of DnaA interaction with repeat variants of *cisII* and *cisMP* elements.

| Repeat No. | Dissociation constants (Kd) [Mean ±SD, μM] | |
| --- | --- | --- |
|  | *cisII* variants | *cisMP* variants |
| Full length | 0.31 ± 0.004 | 0.56 ± 0.21 |
| 8/5 repeats | 0.70 ± 0.35 | 0.89 ± 0.39 |
| 3 repeats | 0.80 ± 0.11 | 1.4 ± 0.7 |
| 2 repeats | Poor affinity | Poor affinity |
| 1 repeat | No affinity | No affinity |

### 3.4 *cisII* and *cisMP* elements contain sites for specific interaction of ParB proteins

Earlier, we had shown the sequence specific interaction of ParB2 and ParB3 with both *cisII* and *cisMP albeit* with different affinity^36^. Since, ParBs function in genome segregation as dimer, we wanted to check the minimum number of repeats required for their sequence specific interaction with cognate ParBs. The radiolabelled *cisII* and *cisMP* and their repeat variants were incubated with purified recombinant ParB2 and ParB3 proteins (Fig. S1) and products were resolved on native PAGE. The ParB2 and ParB3 showed higher affinity with full length *cisII* (Kd = 0.40 ± 0.007 μM) and *cisMP* (0.59 ± 0.05 μM), respectively (Fig. 4 A, E). The affinity of ParBs to cognate *cis* elements were nearly same upto 3 repeats and reduced significantly when the number of repeats become less than 3 (Fig. 4C, G; Table 1). These proteins could rarely bind when number of repeats were less than 2 (Fig. 4 D, H). The ParBs binding to cognate *cis* elements was sequence specific as the ParB-*cis* nucleoprotein complex was not affected in the presence of a 100-fold higher molar concentration of non-specific DNA (Fig 4 A-C, E-G). Further, it was noticed that the full length cisII and cisMP bound ParBs migrated under electrophoresis with the peculiar patterns that normally seen when a tertiary or quaternary structure is introduced either naked DNA or by binding with proteins. Since, reduction in number of repeats has abolished this pattern, a possibility of full-length cis elements producing structures upon protein binding cannot be ruled out. Nonetheless, these findings suggested that both *cisII* and *cisMP* elements have specific sites for ParB nucleation, which might function as centromere - like sequence during segregation of cognate replicons.

**Figure 4.**
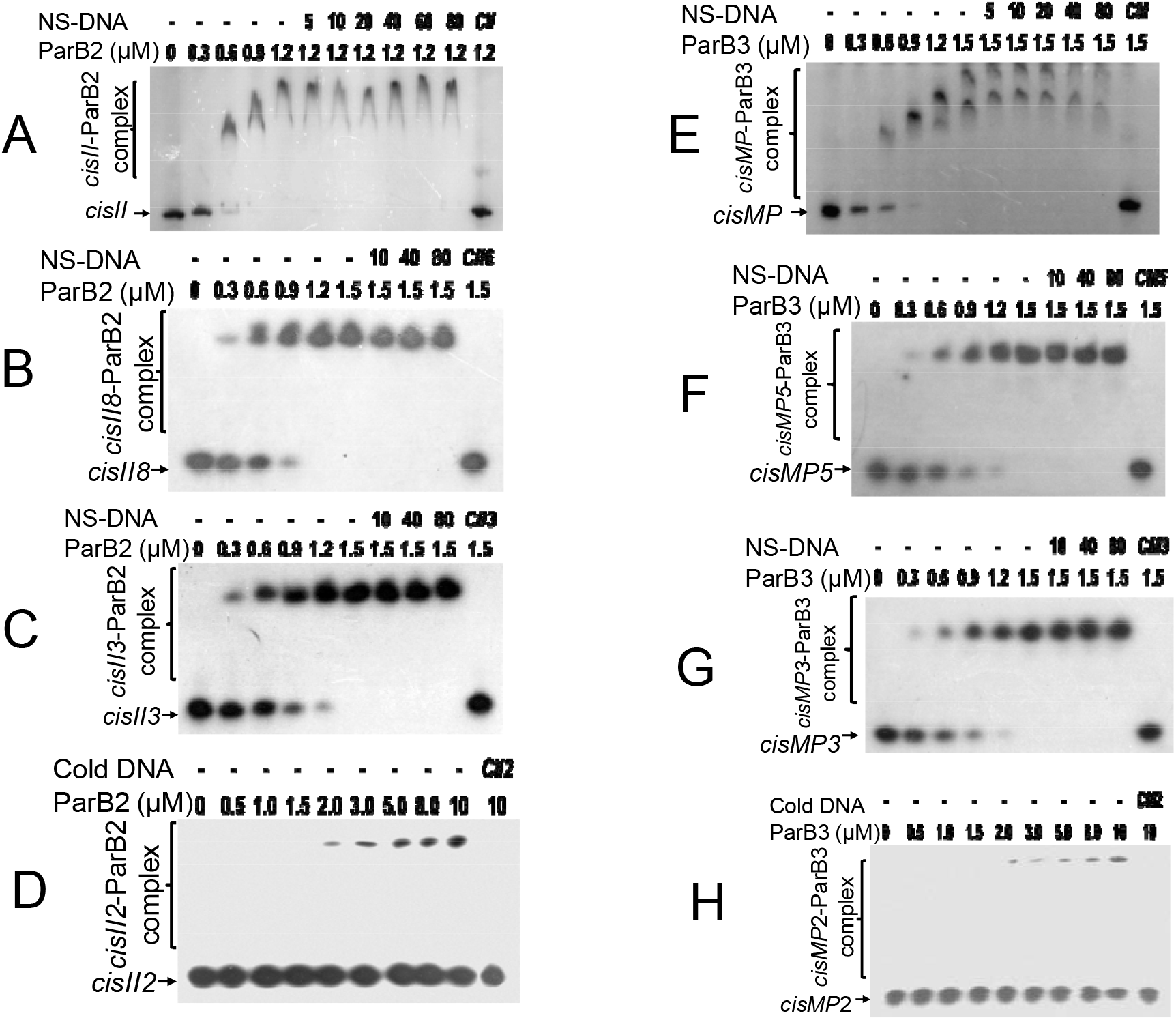
ParBs interaction with *cisII* and *cisMP* and their repeats variants. The radiolabelled full length *cisII* containing 11 repeats (A), 8 repeats (B), 3 repeats (C) and 2 repeat (D) (Table S2B) were incubated with the increasing concentration of ParB2. Similarly, the radiolabelled full length *cisMP* containing 8 repeats (E), 5 repeats (F), 3 repeats (G) and 2 repeats (H) were incubated with the increasing concentration of ParB3. A saturating concentration of DNA-protein ratio was chased with increasing molar ratio of non-specific DNA (NS-DNA) and products were analysed on native PAGE. The fraction of DNA bound with proteins were quantified densitometrically and plotted as a function of protein concentration using Graphpad PRISM. The Kd for the curve fitting of individual plots was determined and given in Table 2.

**Table 2.** The dissociation constant (Kd) value of ParB2 and ParB3 interaction with repeats variants of *cisII* and *cisMP* elements, respectively.

| No. of repeats | Dissociation constants (Kd) [Mean ±SD, μM] | |
| --- | --- | --- |
|  | ParB2 with *cisII* variants | ParB3 with *cisMP* variants |
| Full length | 0.40 ± 0.006 | 0.59 ± 0.06 |
| 8 / 5 repeats | 0.68 ± 0.06 | 0.81 ± 0.04 |
| 3 repeats | 1.06 ± 0.26 | 1.01 ± 0.15 |
| 2 repeats | Poor affinity | Poor affinity |
| 1 repeat | No affinity | Poor affinity |

### 3.5 *cisII* and *cisMP* determine cognate replicons copy number and their segregation

Since both *cisII* and *cisMP* have *‘ori’* signatures, their roles in the regulation of genome copy number and their segregation pattern were investigated. For that, the deletion mutants of *cisII* (Δ*cisII*) and *cisMP* (Δ*cisMP*) elements were made in *D. radiodurans* (Fig. S3) and genome copy number in these mutants was determined by qPCR. The copy number of Chr II and MP was drastically reduced in Δ*cisII* and Δ*cisMP* cells, respectively while there was no significant change for Chr I (Fig. 5A). For instance, the average copy number of Chr II reduced from 5.95±0.32 to 1.55 ±0.22 while the copy number of MP showed an insignificant change from 10.85±0.24 to 8.52±0.81 in Δ*cisII* mutant. Similarly, the average copy number of MP was reduced from 10.85±0.24 to 2.42±0.35 in Δ*cisMP* mutant (Fig. 5 A). Deletion of *cisMP* has not affected the copy number of Chr I and Chr II when compared with wild type. Such a reduction in ploidy of secondary genome elements after deletion of their cognate *cis* elements could be due to either defective replication and/or segregation of these replicons.

**Figure 5.**
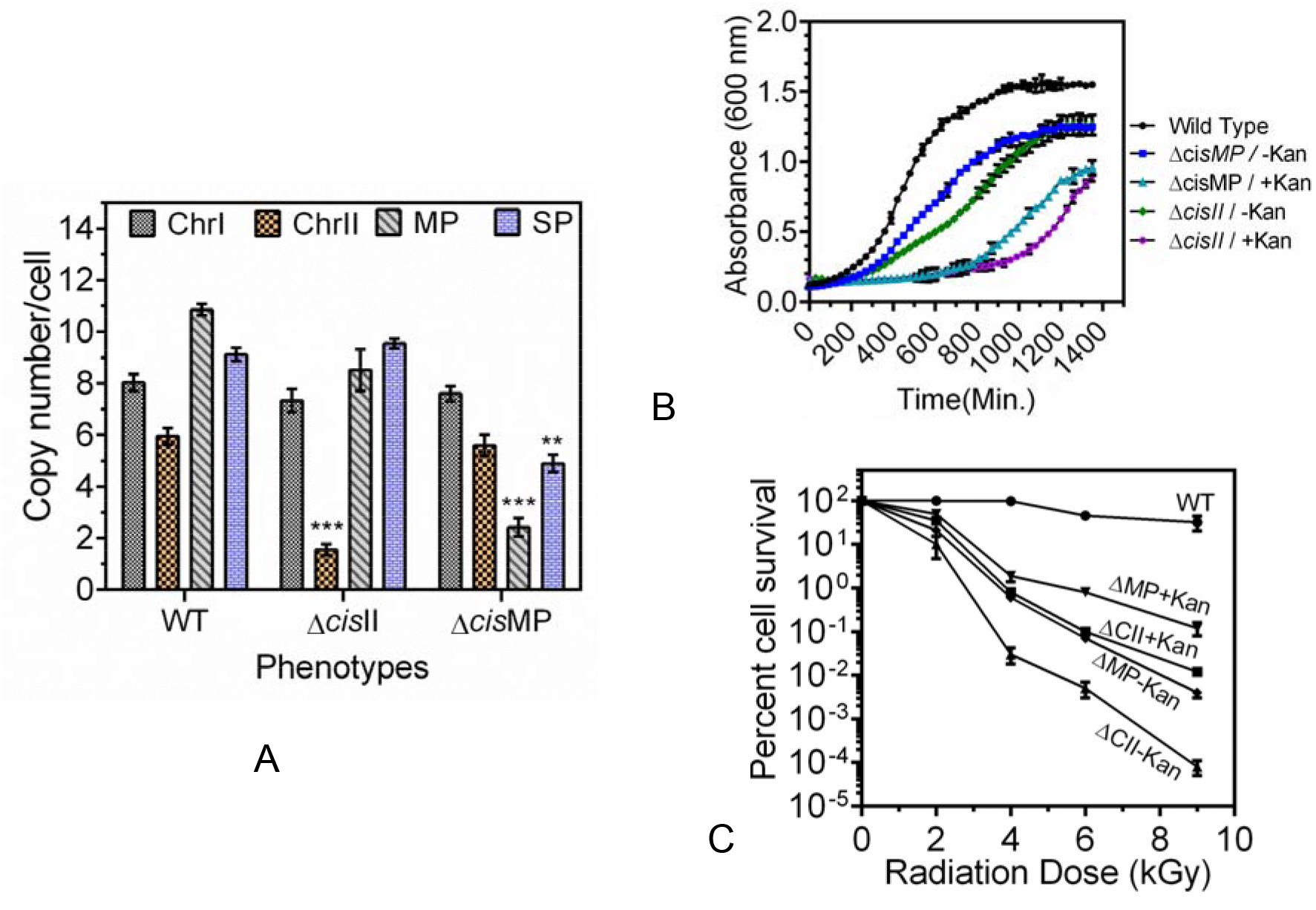
Effect of *cisII* and *cisMP* deletion on genome copy number and radioresistance in *D. radiodurans*. The exponentially growing *D. radiodurans* R1 (WT) and its *cisII* (△cisII) and *cisMP* (△cisMP) deletion mutants were used for the determination of copy number of each replicon (A). Both wild type and *cis* mutants of *D. radiodurans* were checked for growth in the presence and absence of kanamycin under normal conditions (B). Similarly, the gamma radiation response of wild type (WT) and its *cisII* (△CII) and *cisMP* (△MP) mutants grown in the presence (+Kan) and absence (-Kan) of kanamycin was monitored (C).

The possibility of Δ*cisII* and Δ*cisMP* mutants have defects in segregation was checked indirectly by monitoring the frequency of cells conferring antibiotic resistance (integrated in Chr II and MP) when grown in the absence and presence of selection pressure. In general, the growth pattern of *cis* mutants was different from wild type. Further, the growth of both the mutants had reduced when grew in the presence of antibiotics as compared to the absence of antibiotics (Fig. 5 B). Since, antibiotic is being expressed in place of *cis* elements in respective replicons, the differences in the number of cells in the absence versus presence of antibiotic could be an indication of segregation defect in dividing population and secondary genome elements are dispensable for normal growth of this bacterium. Similar findings have been reported earlier in *Bacillus subtilis* and *Mycobacterium tuberculosis* ^46–49^. These results together suggested that *cisII* and *cisMP* elements carry functions for controlling both DNA replication and segregation in *D. radiodurans*.

### 3.6 *cisII* and *cisMP* mutants showed reduction in γ-radiation resistance

Since, the deletion of *cisII* and *cisMP* in *D. radiodurans* has drastically reduced the copy number of respective replicons, a possibility of this if has affected the extraordinary resistance to DNA damage in this bacterium was hypothesized and tested. We monitored the survival of Δ*cisII* and Δ*cisMP* mutants under γ-radiation exposure and found that Δ*cisII* mutant was in general more sensitive to gamma radiation than Δ*cisMP* mutant. Further, the mutants that were maintained without antibiotics showed nearly 2-fold higher sensitivity to gamma radiation as compared to those grown with antibiotics (Fig 5C). Since, the deletion of these *cis* elements had caused reduction in cognate replicon copy number, the loss of gamma radiation resistance could be due to either reduction in copy number or complete loss of secondary genome elements in these mutants. Earlier, it was shown that the reduction of copy number of Chr II and MP in △*parA2*△*parA3* double mutants has affected gamma radiation response in *D. radiodurans* ^36^. These results suggested that deletion of *cisII* or *cisMP* from genome can make these cells defective in DNA replication and /or segregation of secondary genome elements like Chr II and MP, which might have contributed to the loss radioresistance in this bacterium.

### 3.7 *cisII* and *cisMP* deletions affected the maintenance of cognate genome elements

The multipartite genome system, ploidy and packaging of complete genetic materials into a doughnut shaped toroidal nucleiod are the remarkable cytogenetic feature in *D. radiodurans* ^50–52^. However, the presence of all the multipartite genome elements in the nucleoid has not been physically demonstrated. In this work, we developed **F**luorescent **R**eporter-**O**perator **S**ystem (FROS) approach to study the localization of ChrI, Chr II & MP in wild type and the Δ*cisII* and Δ*cisMP* mutants. All three elements (Chr I, Chr II and MP) were tagged with 240 repeats of *tetO* cassette (Table S1) in wild type and *cis* mutants background (Fig. S4) and TetR-GFP was expressed *in trans* (Fig S5). These cells were grown exponentially and stained with Nile red and DAPI, and imaged microscopically in DIC, FITC, TRITC and DAPI channels. Interestingly, we could not find any change in dynamics and localisation of Chr I in Δ*cisII* and Δ*cisMP* mutant when compared with wild type (Fig. 6). The average number of foci for Chr I per cell was ~3, which were localized away from septum and seems to be radially distributed throughout the cytoplasm in both wild type and Δ*cisII* and Δ*cisMP* mutants. A nearly similar pattern of Chr I localisation has been recently demonstrated in *D. radiodurans*, where number of foci per nucleoid was shown to be in the range of 2.25 to 4.44 ^28^. However, the average number of foci for Chr II and MP were found to be 2.09 and 1.1 per cell and were located away from septum and spanned throughout the nucleoid in wild type. However, the localization pattern of Chr II and MP was different in respective *cis* mutant as compared to wild type. For instance, Chr II localization in nucleoid was altered in Δ*cisII* but not in Δ*cisMP* and vice versa. Number of foci of Chr II had reduced from 2 in wild type to 1 in Δ*cisII* mutant. Both these *cis* mutants produced a population of cells devoid of cognate replicons (Fig. 7, Fig 8). Since, qPCR study has shown highest number (10.85) of copies of megaplasmid per cell, the single GFP foci observed through FROS is intriguing but informative. The similar results have been reported in other limited copy plasmids like RK2, R1 etc. in *E. coli* ^53–54^. The decreasing number of GFP foci from 3-4 in Chr I to 1 in MP agreed with their size difference and if that has affected resolution cannot be ruled out. These results highlighted the roles of *cisII* and *cisMP* as ‘*ori*’ and centromere-like functions in the secondary genome elements of *D. radiodurans.* Further, the FROS based localization provided first evidence of both primary and secondary genome elements as part of compact nucleoid and the independent mechanisms for maintenance of primary chromosome and secondary genome elements in this multipartite genome harboring bacterium.

**Figure 6.**
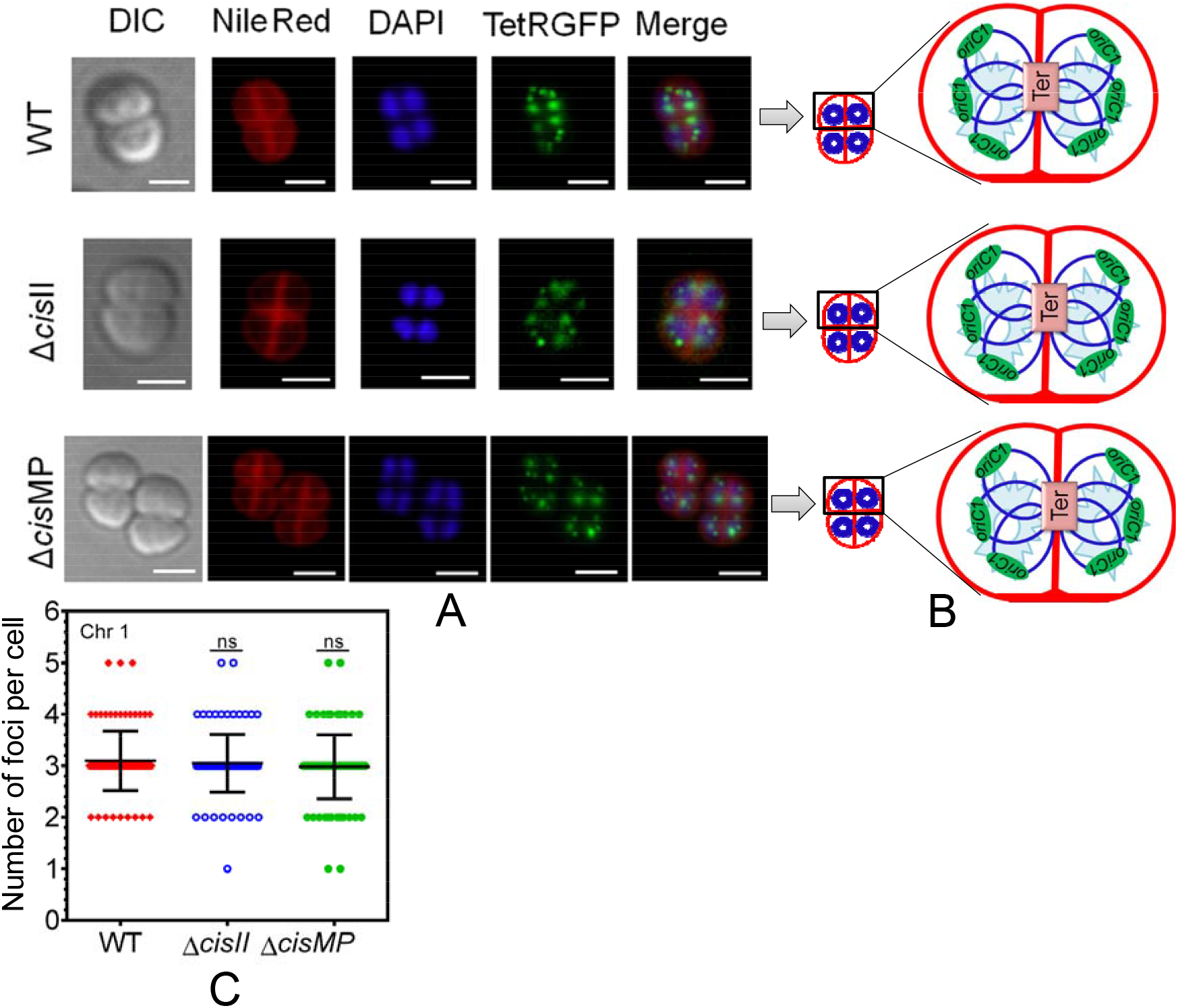
Localization pattern of chromosome I in the nucleiod of wild type and secondary genome *cis* mutants of *Deinococcus radiodurans*. Chromosome I was tagged with tetO/TetR-GFP based FROS system in both wild type (WT) and *cis* mutants *(△cisII and △cisMP)* as described in methods and grown exponentially. These cells were stained with Nile Red and DAPI and visualized microscopically in DIC, TRITC (Nile Red), DAPI (DAPI), FITC (GFP) channels and images were merged (Merge) (A). The schematic diagrams showing the foci position with respect to nucleoid and septum are presented for better clarity (B). Data shown is from a single tetrad where most of these cells show similar pattern as quantified from ~200 cells (C).

**Figure 7.**
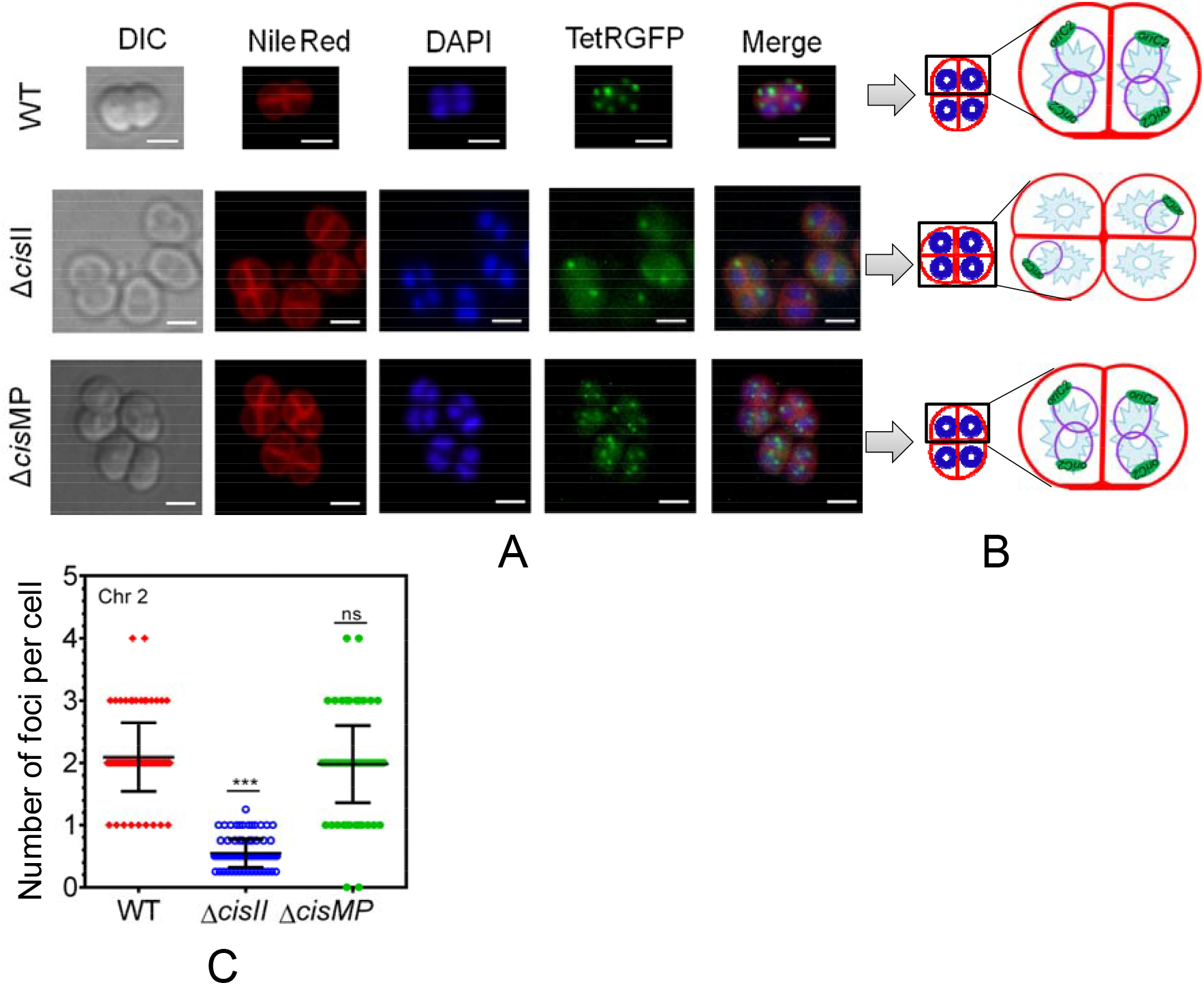
Localization pattern of chromosome II in the nucleiod of wild type and secondary genome *cis* mutants of *Deinococcus radiodurans*. Chromosome II was tagged with tetO-TetR-GFP based FROS system in both wild type (WT) and *cis* mutants (△cisII and △cisMP) as described in methods and grown exponentially. These cells were stained with Nile Red and DAPI and visualized microscopically in DIC, TRITC (Nile Red), DAPI (DAPI), FITC (GFP) channels and images were merged (Merge) (A). The schematic diagrams showing the foci position with respect to nucleoid and septum are presented for better clarity (B). Data shown is from a single tetrad where most of these cells show similar pattern as quantified from ~200 cells (C).

**Figure 8.**
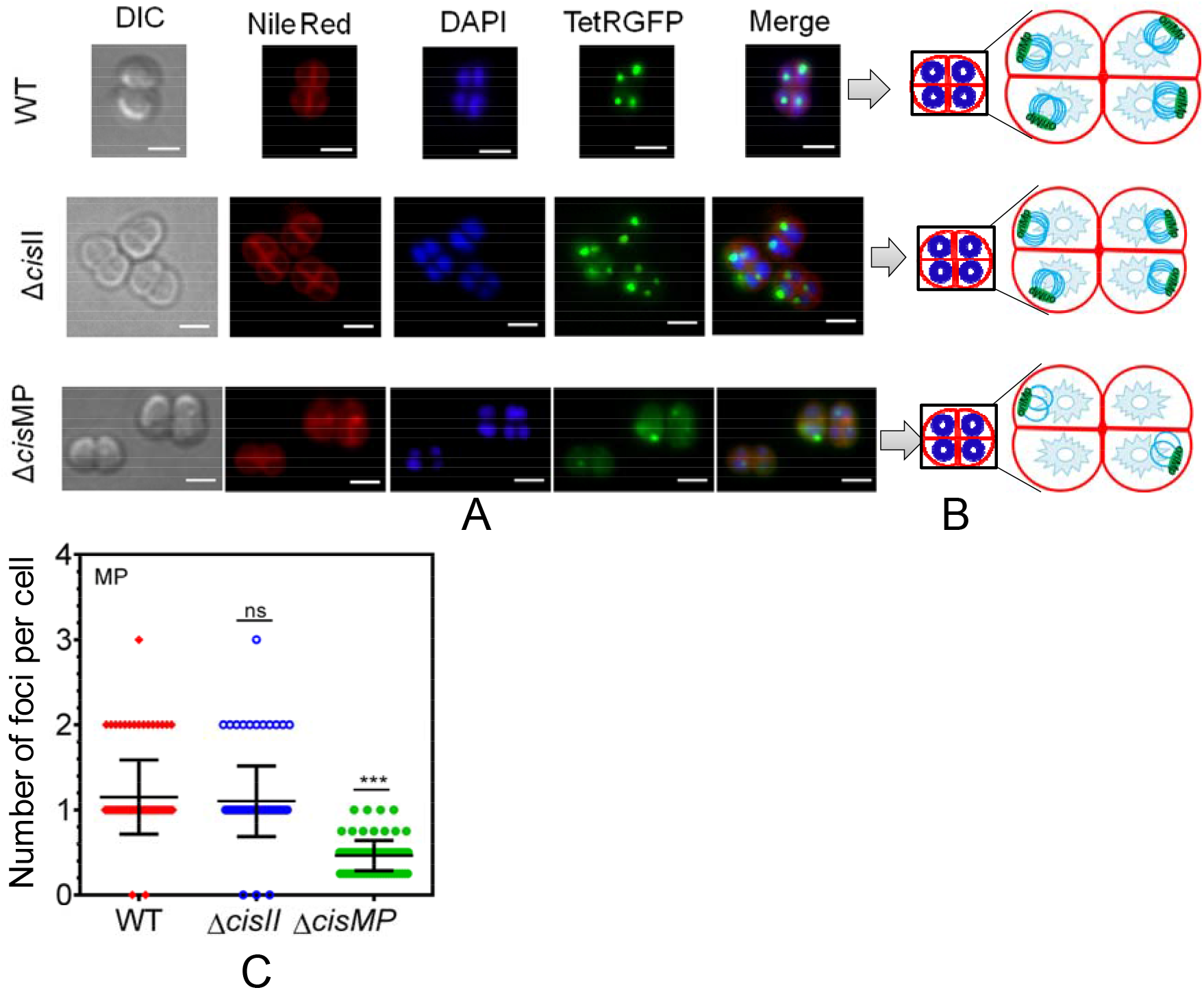
Localization pattern of megaplasmid in the nucleiod of wild type and secondary genome *cis* mutants of *Deinococcus radiodurans*. The megaplasmid was tagged with tetO-TetR-GFP based FROS system in both wild type (WT) and *cis* mutants (△cisII and △cisMP) as described in methods and grown exponentially. These cells were stained with Nile Red and DAPI and visualized microscopically in DIC, TRITC (Nile Red), DAPI (DAPI), FITC (GFP) channels and images were merged (Merge) (A). The schematic diagrams showing the foci position with respect to nucleoid and septum are presented for better clarity (B). Data shown is from a single tetrad where most of these cells show similar pattern as quantified from ~200 cells (C).

## 4. Discussion

The interdependent regulation of DNA replication and segregation process has been studied in bacteria harbouring single circular chromosome. A limited information on these aspects are available in some multipartite genome harbouring bacteria like *V. cholerae, R. sphaeroides, S. meliloti* and *D. radiodurans* ^22, 24, 27, 36, 38^. Though, the nucleoid dynamics and radial positioning of *ori* and *ter* in Chr I of *D. radiodurans* have been demonstrated as a function of cell cycle ^28^, there is not much information available on the organization of the secondary genome elements in this bacterium. This study has brought forth the functional characterization of direct repeats present upstream to *parAB* operon in secondary genome elements; Chr II (*cisII*) and MP (*cisMP*) as ‘*ori*’ and centromere -like sequence in *D. radiodurans,* and used FROS approach to demonstrate the localisation of Chr I, Chr II and MP within the nucleoid in wild type as well as in *cis II* and *cisMP* mutants. We showed that both DnaA and ParBs establish specific interaction with the overlapping *cis* elements indicating the interdependent regulation of both the macromolecular events in *D. radiodurans*. The possibility of such type organization of ‘*ori*’ and centromere-like region is helpful in temporal regulation of replication and segregation in this bacterium. Functionally, the conversion of a non-replicative plasmid into replicative one by these *cis* elements (Fig. 2), reduction in copy number of cognate replicon without these elements (Fig. 5), and increased frequency of Chr II and MP anucleate cells in deletion mutant of cognate *cis* element (Fig. 7, Fig. 8) together support the existence of the functional *‘ori’* and centromere-like sequences within these *cis* elements. The possibility of such type of organization of *cis* elements provide interdependent regulation in two important macromolecular events in bacteria growth cannot be ruled out. Earlier, genome wide studies has shown that most bacteria have centromere-like sequences in close proximity to ‘*oriC*’ in their chromosomes ^12^. Additionally, the molecular crosstalk between the components of these macromolecular events has also been reported in some bacteria. For instance, in *B. subtilis,* Soj (a ParA homologue) directly interacts with DnaA proteins to regulate the replication both positively and negatively at *oriC*, respectively^55^. In *C. crescentus,* DnaA controls chromosome segregation in ParA dependent manner as well as by binding directly to the *parS* site ^11^.

Amongst bacteria harbouring multipartite genome system although, both the chromosomes in *V. cholerae* independently encodes replication and segregation components^56^, the interdependent regulation of these two processes have been reported in this bacterium. For instance, ParA1 a genome segregation protein, encoded on Chr I in *V. cholerae* does stimulate DNA replication through its direct interaction with DnaA while ParB2 (another segregation protein) inhibited the replication initiation in Chr I by yet uncharacterized mechanism^57^. On contrary, ParB2 stimulates replication of Chr II by decreasing RctB binding to inhibitory DNA sequences adjacent to the *oriII* in *V. cholerae* ^58,59^. Existence of both *ori* and centromere-like sequences in close vicinity might offer a fine regulation of these processes by competitive binding of corresponding *trans* factors in *D. radiodurans*. We demonstrated that DnaA can bind specifically to full length of both *cisII* and *cisMP* elements *albeit* different affinities and its affinity have further decreased when the number of repeats has decreased. The loss of DnaA affinity to *cis* elements having lesser number of direct repeats further supports the <u>‘ori’</u> nature of these *cis* elements. Loss of DnaA interaction with *ori*C having lesser number of DnaA boxes has been reported in *M. tuberculosis*, *T. thermophilus* and *H. pylori* ^60–62^. It is noteworthy that *cis* mutants continue to have certain copies of their cognate replicons. This may support a possibility of the *oriC*-independent replication initiation of these replicons, as previously reported in other bacteria ^63^.

Likewise, if these *cis* elements are believed to contain centromere-like sequences, then their deletion should show segregation defect in at least respective replicons. Our results supported the hypothesis and the mutants maintained without antibiotic selection had a heterogenous population of cells with and without cognate replicon as seen where nearly half of the population was sensitive to antibiotics (Fig 5B). Presumably, these were the ones that survived with chromosome I and without secondary genome elements due to segregation defects. Additionally, 2-3 cells in tetrad were devoid of FROS foci in respective genome elements (Fig 7 & Fig 8). When these *cis* mutants were checked for gamma radiation response, the mutants maintained without selection pressure showed a greater sensitivity to gamma radiation as compared to those maintained under selection pressure. This further supported that the cells maintained without selection pressure were heterogenous with respect to secondary genome elements and those without secondary genome would have become sensitive to gamma radiation. Reduction in ploidy of secondary genome elements in *D. radiodurans* was previously found to be associated with reduction in growth and sensitivity for abiotic stresses including γ-radiation^36^. These findings clearly suggest a regulated crosstalk between DNA replication and segregation in bacteria that have evolved a complex multipartite genome system. Earlier, physical interaction of ParBs with DnaA and DnaB proteins have been reported in *D. radiodurans* ^36^. Further, a strong possibility to the proximal existence of *cis* elements involved in DNA replication and segregation, and their temporal regulation through mutual interaction of contemporary *trans* acting factors of these processes could be suggested in *D. radiodurans*.

The multipartite genome with ploidy and toroidal shape of the nucleoid are the most interesting cytogenetic features of this extremophile and in many bacteria that are parasite to some forms of life ^21^. In our opinion, this area of bacterial genome biology offers a great potential with a possible revisit of several paradigms set by working with bacteria harbouring single circular chromosome. Thus, it remains a subject of curiosity that how different replicons are localised in toroidal nucleiod, and if they undergo dynamic change during DNA replication, segregation and DSB repair under normal radiation stressed conditions? Here, we employed tetO-TetR-GFP FROS system and localized Chr I, Chr II and MP in nucleoid of wild type and *cis* mutants during exponential growth. We observed that Chr I as judged with its *ori* was radially distributed in both wild type and *cisII* and *cisMP* mutants. It may be noted that there was no change in ploidy status of Chr I in *cis* mutants. On the other hand, the ~2 foci per cell of Chr II and 1 foci for MP as seen on the nucleiod in all cells of the wild type were clearly altered in the absence of *cisII* and *cisMP*, respectively. The higher frequency of Chr II and MP anucleated cells was observed in △*cisII* and △*cisMP* mutants, respectively. Our results further suggested that different replicons are packaged in single nucleoid and the *‘ori’* regions of these replicons are excluded from the membrane proximal region of the cell as previously reported in other bacteria ^28, 64–66^. It would be worth mentioning that ParAs encoded on Chr II and MP were found to be functionally redundant ^36^, while ParBs were not ^37^, and now *cis* elements are also found to be independently regulating both replication and segregation of these elements.

Although, independent studies are required to understand (i) the dynamics of multipartite genome composition during gamma radiation recovery and normal cell cycle regulation in this bacterium, and (ii) DNA damage responsive territorial change of genome elements in cells recovering from gamma radiation exposure, the available results together conclude that (i) *cisII* and *cisMP* carry both *ori* and centromere-like functions, (ii) primary chromosome (Chr I) and secondary genome elements (Chr II and MP) seems to have independent mechanisms for maintenance, (iii) all the three genome elements like Chr I, Chr II and MP are packaged in toroidal shape nucleoid but seems to be organized differently, (iv) secondary genome elements play important roles in extraordinary radioresistance of this bacterium.

## Acknowledgments

Authors are grateful to Dr Swathi Kota, Dr YS Rajpurohit and Dr Sheetal Uppal for their technical discussion on the work. Ms Shruti Mishra and Ms. Reema Chaudhary for their technical and editorial comments and help in fluorescence imaging. GKM is grateful to Department of Atomic Energy, India for research fellowship.

