## Supplementary material for "Overlapping *oriC* and centromere-like functions in secondary genome replicons determine their maintenance independent of chromosome I in *Deinococcus radiodurans*": Table S1, TableS2, Fig S1-Fig S5

Dr. H. S. Misra

Molecular Biology Division

Bhabha Atomic Research Centre

Mumbai- 400085

^3^ Current address: Assistant Professor, MMV, Banaras Hindu University Varanasi, India – 221005

**Table S1 :** List of oligonucleotides used in this study

| Primer Name | Oligonucleotide Sequences | Purpose/Plasmid |
| --- | --- | --- |
| cisIIFw | 5' GCTCTAGAGATCAAATCTTTTAAAG3' | pNOKcisII, EMSA (cisII; full length) |
| cisIIRw | 5' GCTCTAGACTTAATAGACCTGTAATTG 3' |  |
| cisMPFw | 5’ GCGGGCCC CAAGGACGGCTTCTCTC 3’ | pNOKcisMP, EMSA (cisMP; full length) |
| cisMPRw | 5' CG GAATTC AGTTGCAGACCATAGGGGT 3' |  |
| CIIintFw1 | 5’ TCCCCGCGGTCCAAATG 3’ | EMSA (cisII; 8 repeats) |
| CIIintFw2 | 5’TCCCCGCGGCAAACGGCCCA 3’ | EMSA (cisII; 3 repeats) |
| CIIintFw3 | 5’CTCCACAAAGTGCCACAGGTAATTCCACAAAGTGCCACAGGC 3’ | EMSA (cisII; 2 repeats) |
| CIIintRw3 | 5’GCCTGTGGCACTTTGTGGAATTACCTGTGGCACTTTGTGGAG 3’ |  |
| CIIintFw4 | 5’ CTCCACAAAGTGCCACAGGTG 3’ | EMSA (cisII; 1 repeat) |
| CIIintRw4 | 5’ CACCTGTGGCACTTTGTGGAG 3’ |  |
| CMintFw1 | 5’ GCGGGCCCTTTTGCACGTTG 3’ | EMSA (cisMP; 5 repeats) |
| CMintFw2 | 5’ GCGGGCCCGTCTACAAAGAG 3’ | EMSA (cisMP; 3 repeats) |
| CMintFw3 | 5’ACGCAAAGGTGTCGCTATTTTGACCCCAAATCCCGCAAAGGTGTCGCTAT 3’ | EMSA (cisMP; 2 repeats) |
| CMintRw3 | 5’ATAGCGACACCTTTGCGGGATTTGGGGTCAAAATAGCGACACCTTTGCGT 3’ |  |
| CMintFw4 | 5’ CCCGCAAAGGTGTCGCTAGG 3’ | EMSA (cisMP; 1 repeat) |
| CMintRw4 | 5’ CCTAGCGACACCTTTGCGGG 3’ |  |
| CIIUPFw. | 5’ GGGGTACC TCGGTCACGTCGTATGC 3’ | pNOKCII |
| CIIUPRw | 5’ CGGAATTC CCTATGATGATGATCATC 3’ |  |
| CIIDNFw | 5’ CGGGATCC TTTGTGCTGAAGAATCATC 3’ |  |
| CIIDNRw | 5’ GCTCTAGA AAGGCTAGGCGGACTATC 3’ |  |
| CMPUPFw | 5’ GGGGTACCGACAGAAGTCTTACGGCC 3’ | pNOKCMP |
| CMPUPRw | 5’ CGGAATTCAGCGACACCTTTGCGGGA 3’ |  |
| CMPDNFw | 5’ CGGGATCCCTTGTAAAATTCACCAAC 3’ |  |
| CMPDNRw | 5’ GCTCTAGAGCCCGAGAGAAGGGGGAC 3’ |  |
| pETDAF | 5’ CG GGATCCGTGCGCAAAAACGTCTC 3’ | pETDnaA |
| pETDAR | 5’ GC GAATTCTTACGCCCCGACTTCTTC 3’ |  |
| RTZ Fw | 5' ATCAAGGAATATCTCGA 3' | Quantitative PCR study |
| RTZ Rw | 5' CAGCTTTTCGTTGTTCACC 3' |  |
| RTEFw | 5’ TTGAGCGTCCCCGCGCC3’ |  |
| RTERw | 5’GTGCCCGACGAGGTAGA3’ |  |
| RTKFw | 5’GTGCGGCGCCTGCAACGC3’ |  |
| RTKRw | 5’CCGCCGATAGACAGAATC3’ |  |
| A155RTFw | 5’ TTGGCGCATTTTCCCGGC 3’ |  |
| A155RTRw | 5’ CAGCAGGTTGATCGCCTG 3’ |  |
| PprARTFw | 5’ GTGCTACCCCTGGCCTT 3’ |  |
| PPRARTRW | 5’ GCGGCCATCGGTCAGAAT 3’ |  |
| A182RTFW | 5’ ATTCTGGGGCCGGAGCTG3’ |  |
| A182RTRW | 5’CTTGCGTTCCCCCGGCGG 3’ |  |
| B03RTFw | 5’ CTGAGTCCTGACGAGTCC 3’ |  |
| B03RTRw | 5’ TTCCGGGTGACGCAGCAG 3’ |  |
| B30RTFw | 5’ ATGAGCCGCAAGTTGCCG3’ |  |
| B30RTRw | 5’ GCCAGCGAGGCGGCTCGC 3’ |  |
| B104RTFw | 5’ ATGAAAACTCTTGAGGC3’ |  |
| B104RTRw | `5 GCCGAGGAAAACGTCCAG 3’ |  |
| C01RTFw | 5’ ATGTGCTCGCCTCCTAGA 3’ |  |
| C01RTRw | 5’ TCACTGTGAAACCTGATC 3’ |  |
| C18RTFw | 5’ ATGACACAGACGCGGCG 3’ |  |
| C18RTRw | 5’ GTCCGCGAGGCGCATCAT 3’ |  |
| C34RTFw | 5’TGGTGGCATTTCTCCGTG3’ |  |
| C34RTRw | 5’ CACTGAAATACCCCAGCC3’ |  |
| nptFw | 5’GCACGGTGGCCGAGTGG3’ |  |
| nptmidRw | 5’AACATCATTGGCAACGCT3’ |  |
| blaRTFw | 5’GGATCATGTAACTCGCCT3’ |  |
| blaRTRw | 5’TTACCAATGCTTAATCAGTGAGG3’ |  |
| ChI(1.5 ̊ )Fw | 5’GCTCTAGAGTCGACGCCTCTTTTCACCGCAAAG 3’ | p44Ch1 |
| ChI(1.5 ̊ )Rw | 5’AAAAGTACTCATATGCCGGACATGTCCGGGCGC3’ |  |
| Ch2(4 ̊ )Fw | 5’GCTCTAGAGTCGACGGCAGCGAGGTCAGGAAG 3’ | p44Ch2 |
| Ch2(4 ̊ )Rw | 5’AAAAGTACTAAGCTTACGTCCGGCAAGCACCTG3’ |  |
| MP(4.4 ̊ )Fw | 5’GCTCTAGAGTCGACGAAGCTGGTAAAACTTTG3’ | p44MP |
| MP(4.4 ̊ )Rw | 5’AAAGTACTCATATGTACGCCCGAAAGCCTACAG3’ |  |
| SpecFw | 5’ATGAGGGAAGCGGTGATC3’ | p44SCh1, p44SCh2 and p44SMP construction and Diagnostic PCR |
| SpecRw | 5’TTATTTGCCGACTACCT3’ |  |
| TetRscIApIFw | 5’CGGAGCTCGGGCCCGTGAGATTAGATAAAAG3’ | pDTRGFP & pRADTRGFP |
| TetRSalIRw | 5’GCGTCGACAGACCCACTTTCACATTTAAG3’ |  |
| GFPXbaIRw | 5’GCTCTAGATTATTTGTATAGTTCATCCA3’ |  |
| AmpRw | 5’TTACCAATGCTTAATCAGTGAGG3’ | Diagnostic PCR |
| Dr0010F | 5’GTGAAATCACCGCTTCCAATG3’ |  |
| DrA005F | 5’ATGAAAGCAATTGTCTGGCAAG3’ |  |
| DrB003F | 5’TTGGCCCTGCAACCGGAAC3’ |  |
| A2AfIIIFw | 5’CCCACGTGTTCTGCGCCGCGCTTTAGC 3’ | Primers for non-specific DNA in EMSA with drDnaA/ParBs |
| A2SxIRw | 5’GTACGACCTGGTCACGTCGGCATGAACGGGT 3’ |  |
| 260bpFw | 5'ATCAAGGAATATCTCGA3' (RTZFW) |  |
| 260bpRw | 5'CAGCTTTTCGTTGTTCACC3' (RTZRW) |  |
| 100bpFw | 5'ATCAAGGAATATCTCGA3' (RTZFW) |  |
| 100bpRw | 5’GGGCAATTTCGGCCACGAC3’ |  |

**Table S2: List of bacterial strains and plasmids used in this study**

| **Bacterial strains** | | **Genotype** | | | **Source** | |
| --- | --- | --- | --- | --- | --- | --- |
| *D. radiodurans* R1 | | Wild type strain ATCC13939 | | | Lab stock | |
| *ΔcisII* | | *cisII* (region; 150-515) replaced with *nptII* cassetess from chromosome II of *D. radiodurans* R1 (Kan^R^) | | | This study | |
| *ΔcisMP* | | *cisMP* (region; 177403- 400) replaced with *nptII* cassetess from megaplasmid of *D. radiodurans* R1 ( Kan^R^) | | | This study | |
| R1::ChrI-*tetO* | | p44SCh1 plasmid integrated at 1.5˚ position in chromosome I of wild type (Spec^R^) | | | This study | |
| Δ*cisII*::ChrI-*tetO* | | p44SCh1 plasmid integrated at 1.5˚ position in chromosome I of *ΔcisII* (Kan^R^; Spec^R^) | | | This study | |
| Δ*cisMP*::ChrI-*tetO* | | p44SCh1 plasmid integrated at 1.5˚ position in chromosome I of *ΔcisMP* (Kan^R^; Spec^R^) | | | This study | |
| R1::ChrII-*tetO* | | p44SCh2 plasmid integrated at 4˚ position in chromosome II of wild type (Spec^R^) | | | This study | |
| Δ*cisII*::ChrII-*tetO* | | p44SCh2 plasmid integrated at 4˚ position in chromosome II of *ΔcisII* (Kan^R^; Spec^R^) | | | This study | |
| Δ*cisMP*::ChrII-*tetO* | | p44SCh2 plasmid integrated at 4˚ position in chromosome II of *ΔcisMP* (Kan^R^; Spec^R^) | | | This study | |
| R1::MP-*tetO* | | p44SMP plasmid integrated at 4.4˚ position in megaplasmid of wild type (Spec^R^) | | | This study | |
| Δ*cisII*::MP-*tetO* | | p44SMP plasmid integrated at 4.4˚ position in megaplasmid of *ΔcisII* (Kan^R^; Spec^R^) | | | This study | |
| Δ*cisMP*::MP-*tetO* | | p44SMP plasmid integrated at 4.4˚ position in megaplasmid of *ΔcisMP* (Kan^R^; Spec^R^) | | | This study | |
| *ΔrecA* | | Deinococcal *recA* gene disrupted with chloramphenicol resistance gene cassettes (Cam^R^) | | | Lab stock | |
| *E. coli* NovaBlue | | *end*A1 *hsd*R17*(r _K12_ ^−^ m _K12_ ^+^) sup*E44 *thi-1 rec*A1 *gyr*A96 *rel*A1 *lac*F’'*[pro*A^+^B*^+^ lac*I^q^ *Z∆*M15*::*T*n*10 ] (Tet^R^) | | | NEB Inc., | |
| *E. coli* BL21(DE3) | | *fhu*A2 *(lon)omp*T *gal*(λDE3)*(dcm) ∆hs*dS | | | Lab stock | |
| **Plasmids** | | | | | | |
| **Sr No.** | **Plasmids** | | **Characteristics** | **Sources** | | **MW of**  **Protein**  **(~kDa)** |
| 1 | pET28a(+) | | ~ 5.3 kb plasmid; N-terminal 6XHis tag (Kan^R^) | Novagen | | - |
| 2 | pETDnaA | | pET28a(+) carrying Dr_0002 at *Bam*HI and *Eco*RI | This study | | ~ 53 kDa |
|  | pETB2 | | pET28a(+) carrying Dr_A0002 at *Nde*I and *Xho*I | Maurya et al., 2019a | | ~33 kDa |
|  | pETB3 | | pET28a(+) carrying Dr_B0002 at *Nde*I and *Xho*I | Maurya et al., 2019a | | ~32 kDa |
| 3 | pNOKOUT | | A deinoccocal suicidal vector; Kan^R^ | Kahirnar et al., 2008 | | - |
| 4 | pNOKcisII, | | pNOKOUT carrying full length *cisII* at *Xba*I site | This study | | - |
| 5 | pNOKcisMP | | pNOKOUT carrying full length *cisMP* at *Apa*I-*Eco*RI sites | This study | | - |
| 6 | pNOKCII | | pNOKOUT carrying ~500 bps upstream at *Kpn*I & *Eco*RI and ~500 bps downstream at *Bam*HI & *Xba*I of *cisII* element (Kan^R^) | This study | | - |
| 7 | pNOKCMP | | pNOKOUT carrying ~500 bps upstream at *Kpn*I & *Eco*RI and ~500 bps downstream at *Bam*HI & *Xba*I of *cisMP* element (Kan^R^) | This study | | - |
|  | pLAU44 | | Plasmid with an array of 240 repeats of *tetO* (Amp^R^& Gen^R^) | Lau *et al*., 2003 | | - |
|  | p44SCh1 | | pLAU44 with region (10713-11715) from chromosome I at *Xba*I-*Sca*I, with Spectinomycin cassette from pVHS559 at *Nhe*I-*Xho*I | This study | | - |
|  | p44SCh2 | | pLAU44 with region (4695-c5691) from chromosome II at *Xba*I-*Sca*I, with Spectinomycin cassette from pVHS559 at *Nhe*I-*Xho*I | This study | | - |
|  | P44SMP | | pLAU44 with region (2203-3000) from megaplasmid at *Xba*I-*Sca*I, with Spectinomycin cassette from pVHS559 at *Nhe*I-*Xho*I | This study | | - |
|  | pDSW208 | | PDSW208-MCS-*gfp* (fusion vector) (Amp^R^) | Weiss et al., 1999 | | ~27 kDa |
|  | pLAU53 | | pBAD24 backbone containing *lacI*-eCFP and *tetR*-eYFP behind the same *araB* promoter, (Kan^R^ & Amp^R^) | Lau et al., 2003 | | - |
|  | pDTRGFP | | pDSW208 with *tetR* from pLAU53at *Sac*I-*Sal*I to give *gfp-tetR* | This study | | ~48 kDa |
|  | pRADTRGFP | | pRADgro with *gfp-tetR* at *Apa*I-*Xba*I from pDTRGFP | This study | | ~48 kDa |

**Supplementary Figures with legend**

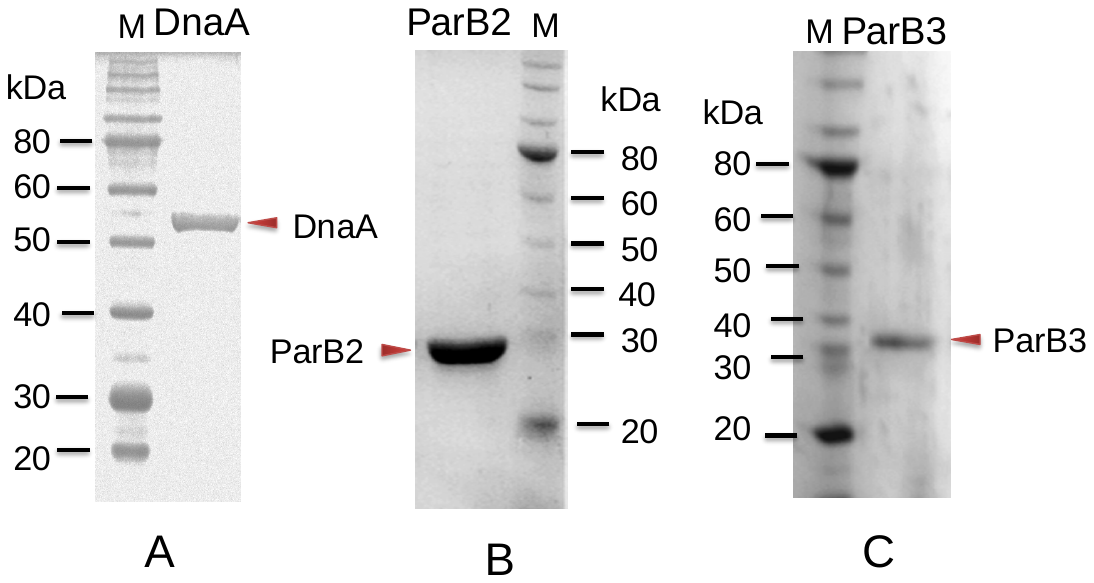

Figure S1. **Purification of recombinant proteins used for DNA protein interaction.** The recombinant DnaA (A), ParB2 (B) and ParB3 (C) were purified from *E. coli* using metal affinity chromatography.

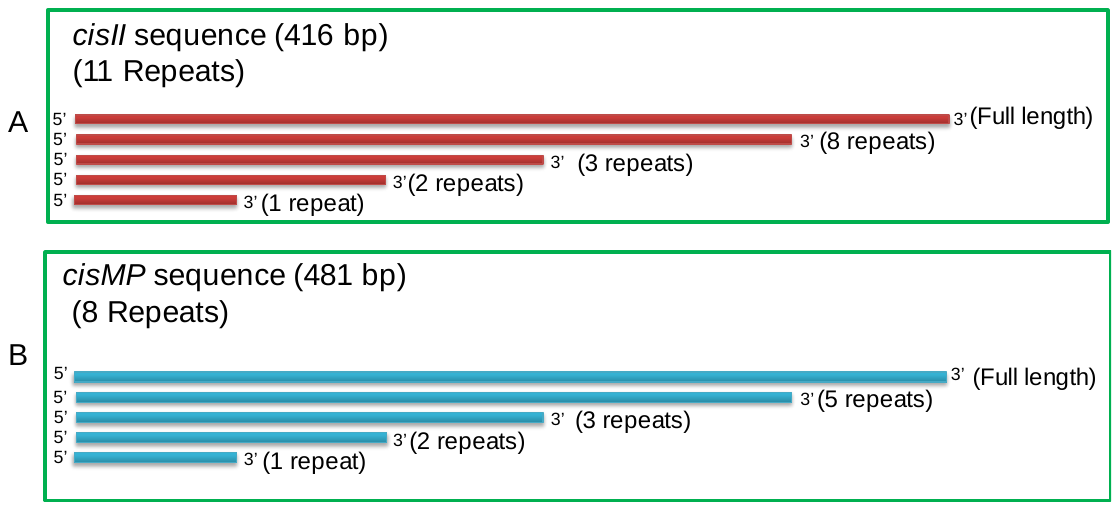

Figure S2. Schematic representation of cisII (A) and cisMP (B) elements and their repeat variants PCR amplified and used for DNA protein interaction studies.

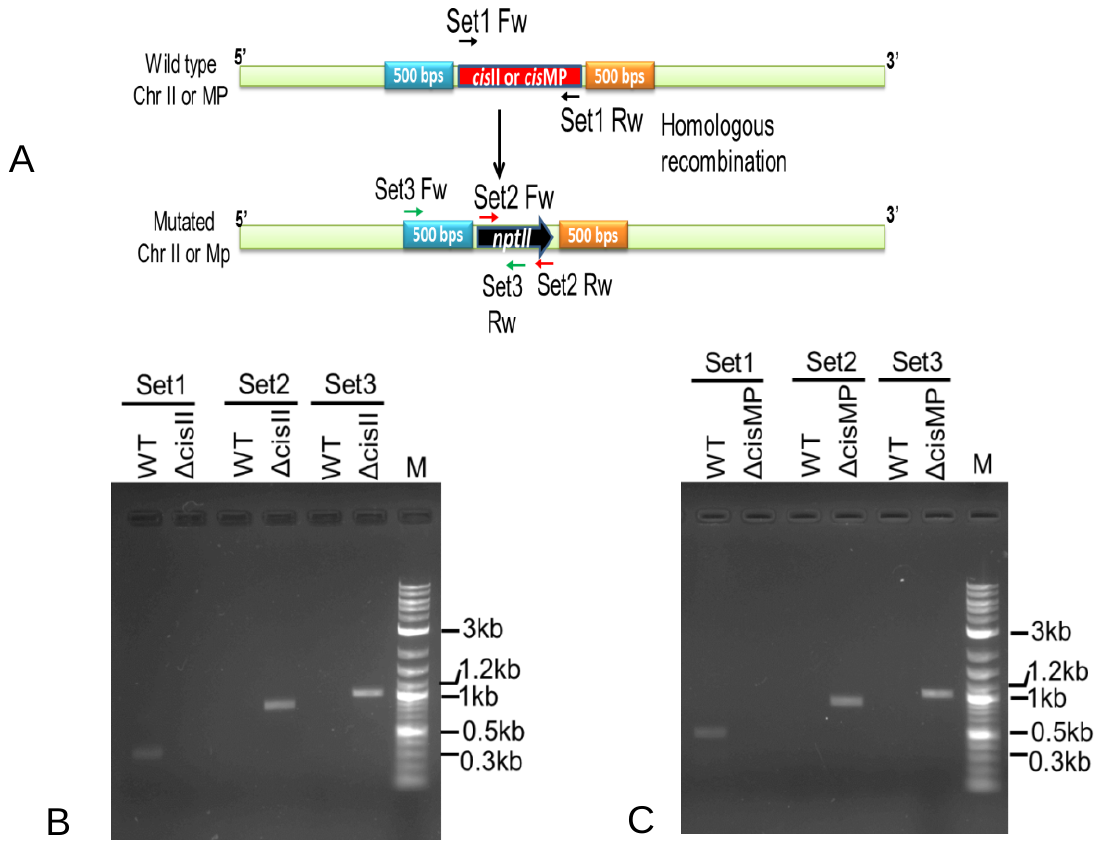

Figure S3**. Generation of *cisII* and *cisMP* deletion mutants in *Deinococcus radiodurans*.** Schematic representation of strategy for replacement of *cis* elements with the expressing cassette of kanamycin and position of primers used for diagnostic PCR (A). The recombinant cells expected to have cisII (B) and cisMP (C) deletions grown several generations under selection pressure. The genomic DNA was prepared, and PCR amplification was carried out by diagnostic PCR using primer as shown in A. Products were analysed on 1% agarose gel.

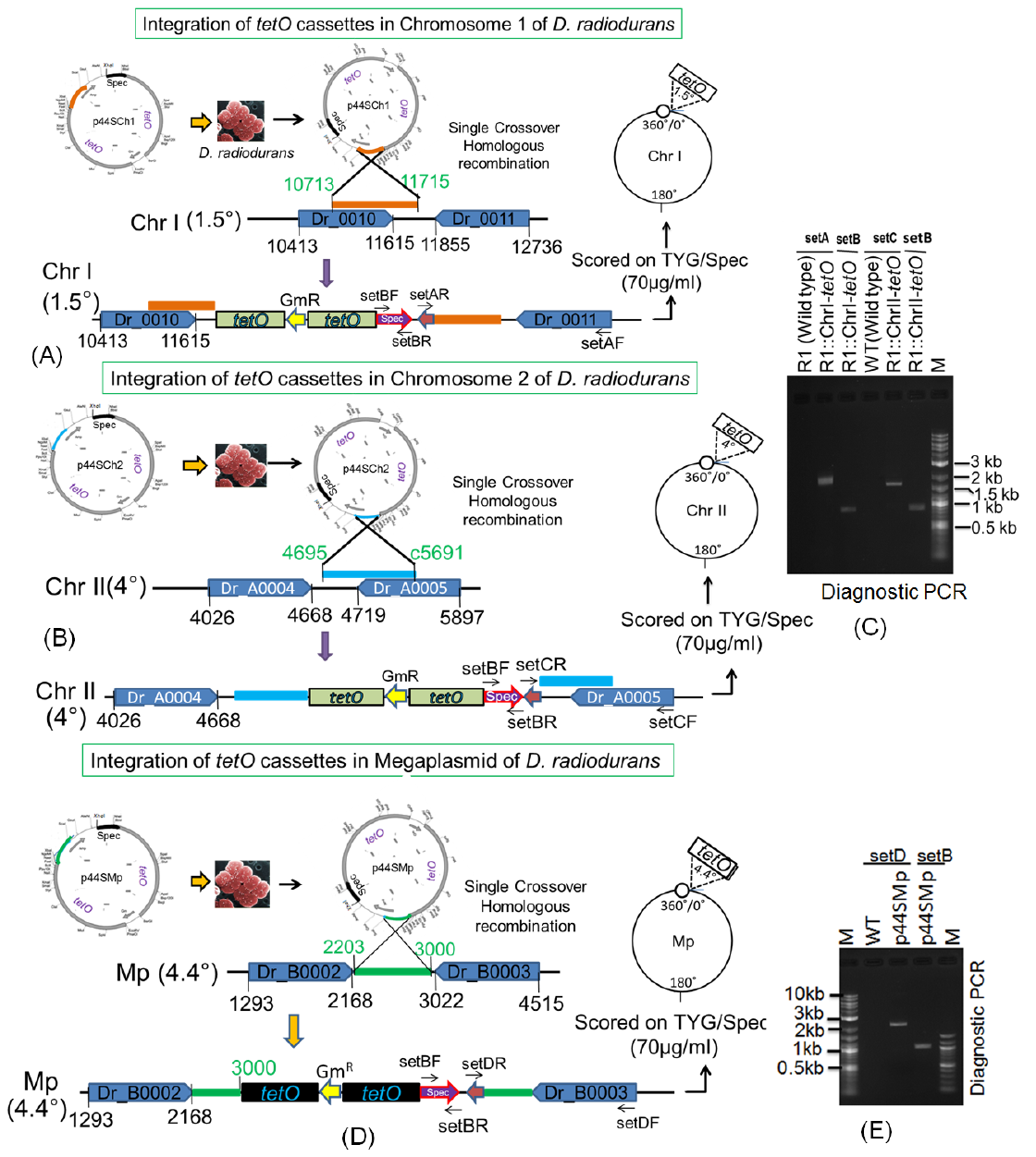

Figure S4. **Schematic representation of tetO-TetR mediated FROS development for D. radiodurans.** The *tetO* cassette from was inserted at 1.5^o^ in chromosome I (A), at 4^o^ in chromosome II (B) and at 4.4^o^ in megaplasmid (D) through homologous recombination. Insertion of *tetO* cassettes into respective positions was confirmed by diagnostic PCR using plasmid sequence specific primers (C and E).

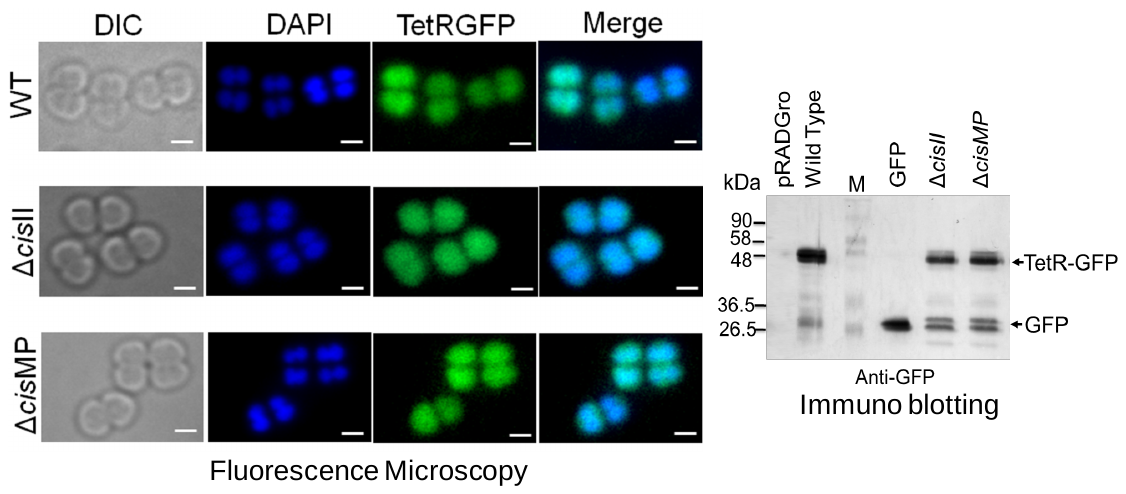

Figure S5. **Expression pattern of TetR-GFP in wild type and cis mutants without *tetO* repeats.** The expression of TetR-GFP in both wild type and cis mutants of *D. radiodurans* was confirmed microscopically (A) and through immunoblotting (B). The TetR-GFP expression in the absence of *tetO* element was found to be uniformly spread in the cell.
